# Characterization of highly proliferative decidual precursor cells during the window of implantation in human endometrium

**DOI:** 10.1101/2020.12.16.423007

**Authors:** Maria Diniz-da-Costa, Chow-Seng Kong, Katherine J. Fishwick, Thomas Rawlings, Paul J. Brighton, Amelia Hawkes, Joshua Odendaal, Siobhan Quenby, Sascha Ott, Emma S Lucas, Pavle Vrljicak, Jan J Brosens

## Abstract

Pregnancy depends on the wholesale transformation of the endometrium, a process driven by differentiation of endometrial stromal cells (EnSC) into specialist decidual cells. Upon embryo implantation, decidual cells impart the tissue plasticity needed to accommodate a rapidly growing conceptus and invading placenta, although the underlying mechanisms are unclear. Here we characterize a discrete population of highly proliferative mesenchymal cells (hPMC) in midluteal human endometrium, coinciding with the window of embryo implantation. Single-cell transcriptomics demonstrated that hPMC express genes involved in chemotaxis and vascular transmigration. Although distinct from resident EnSC, hPMC also express genes encoding pivotal decidual transcription factors and markers, most prominently prolactin. We further show that hPMC are enriched around spiral arterioles, scattered throughout the stroma, and occasionally present in glandular and luminal epithelium. The abundance of hPMC correlated with the *in vitro* colony-forming unit activity of midluteal endometrium and, conversely, clonogenic cells in culture express a gene signature partially conserved in hPMC. Cross-referencing of single-cell RNA-sequencing data sets indicated that hPMC differentiate into a recently discovered decidual subpopulation in early pregnancy. Finally, we demonstrate that recurrent pregnancy loss is associated with hPMC depletion. Collectively, our findings characterize midluteal hPMC as novel decidual precursors that are likely derived from circulating bone marrow-derived mesenchymal stem/stromal cells and integral to decidual plasticity in pregnancy.

**Significance statement:** Transformation of cycling endometrium into the decidua of pregnancy requires extensive tissue remodeling. Perturbations in this process lead to breakdown of the maternal-fetal interface and miscarriage. Here we report the characterization of decidual precursor cells during the window of implantation. We demonstrate that decidual precursors are clonogenic and primed for exponential growth. They likely originate from bone marrow-derived MSC and give rise to a distinct decidual subpopulation in pregnancy. Recurrent pregnancy loss is associated with loss of decidual precursor cells prior to conception, raising the possibility that they can be harnessed for the prevention of pregnancy disorders, including miscarriages and preterm labor.

## Introduction

The uterine mucosa – the endometrium – is a highly regenerative tissue capable of adopting different physiological states during the reproductive years [1,2]. In the midluteal phase of the menstrual cycle, the endometrium starts remodeling intensively in response to elevated progesterone levels and rising intracellular cyclic AMP levels [3], heralding the start of a short window during which an embryo can implant [4]. This process, termed decidualization, begins with an acute stress response in endometrial stromal cells (EnSC) [5], resulting in cell cycle arrest, release of proinflammatory mediators, and influx of uterine natural killer (uNK) cells [6,7]. After a lag period of approximately 4 days, phenotypic decidual cells emerge, coinciding with the closure of the implantation window [6–8]. Decidual cells are progesterone-dependent secretory cells with a well-developed endoplasmic reticulum and enlarged nuclei. They are refractory to various environmental stress signals [9–12], and form a quasi-autonomous, anti-inflammatory matrix around the implanting embryo, which enables invading extravillous trophoblast to form a hemochorial placenta [3]. In addition, the maternal decidua creates a tolerogenic microenvironment that harbors an abundance of infiltrating immune cells, foremost uNK cells but also macrophages, dendritic cells, and regulatory T cells [13].

Recently, we provided evidence that inflammatory reprogramming of EnSC also leads to the emergence of senescent decidual cells [6,8,14]. Unlike decidual cells, senescent decidual cells are progesterone resistant [8]. They secrete a complex mixture of extracellular matrix (ECM) proteins and proteinases, proinflammatory cytokines, and chemokines, designated senescence-associated secretory phenotype (SASP), which causes sterile inflammation and induces secondary senescence in neighboring decidual cells [6,8]. Hence, in a non-conception cycle, falling progesterone levels during the late-luteal phase result in a preponderance of senescent decidual cells, an influx of neutrophils, ECM breakdown, and menstrual shedding of the superficial endometrial layer [15]. By contrast, continuous progesterone signaling upon successful embryo implantation enables decidual cells to engage and activate uNK cells, which in turn target and eliminate senescent decidual cells through perforin- and granzyme-containing granule exocytosis [6,8]. Perturbations that lead to an excessive or persistent pro-senescent decidual response presents the embryo with a uterine mucosa that is easily to invade but also prone to breakdown. Clinically, a pro-senescent decidual response is associated with rapid conceptions (i.e. ‘super-fertility’) and recurrent pregnancy loss [8,16,17].

Despite these emerging insights, it remains unclear how the maternal decidua acquires the necessary plasticity to accommodate rapid expansion in early gestation. Studies in mice have provided compelling evidence that pregnancy is associated with engraftment of circulating bone marrow-derived mesenchymal stem/stromal cells (BDMSC) into the decidua [18–20], which differentiate into nonhematopoietic prolactin-expressing decidual cells [20]. A recent study demonstrated that bone marrow transplants from wild-type mice to mice carrying a heterozygous deletion of *Hoxa11*, a pivotal decidual transcription factor, not only restore the decidual response but also prevent pregnancy loss in these animals [20]. This observation both underscores the importance of BDMSC in establishing a robust placental-decidual interface and signposts a novel strategy for the prevention of intractable pregnancy disorders, such as recurrent pregnancy loss and preterm birth. Whether BDMSC contribute to the decidual transformation of human endometrium is, however, not known.

Based on single-cell transcriptomics, we recently described the presence of a discrete population of highly proliferative mesenchymal cells (hPMC) in midluteal endometrium [8]. Here, we report on the characterization of these cells. Our findings indicate that hPMC are the human equivalent of bone marrow-derived decidual precursor cells in mice. We provide evidence that hPMC impart clonogenicity of midluteal endometrium and give rise to a distinct subpopulation of decidual cells at the maternal-fetal interface in pregnancy. We further demonstrate that a lack of these novel human decidual precursor cells is associated with recurrent pregnancy loss.

## Materials and Methods

### Endometrial sample collection

The collection of endometrial biopsies for research purposes was approved by the NHS National Research Ethics – Hammersmith and Queen Charlotte’s & Chelsea Research Ethics Committee (REC reference: 1997/5065) and Tommy’s National Reproductive Health Biobank (REC reference: 18/WA/0356). Samples were timed relatively to the pre-ovulatory luteinizing hormone (LH) surge and obtained using a Wallach Endocell® endometrial sampler following transvaginal ultrasonography from patients attending a dedicated research clinic at University Hospitals Coventry and Warwickshire (UHCW) National Health Service (NHS) Trust, Coventry, UK. Written informed consent was obtained prior to tissue collection in accordance with the guidelines of the Declaration of Helsinki, 2000. Endometrial biopsies were timed 5 to 10 days after the pre-ovulatory LH surge. Demographic details of the tissue samples used in different experiments are shown in Table S1.

### Endometrial tissue processing, primary cell culture, and CFU assay

Processing of endometrial biopsies, isolation of sushi domain containing 2 (SUSD2)-positive cells using magnetic-activated cell sorting, and culturing of primary endometrial cells were carried out as described in our published step-by-step protocol [21]. CFU assays were also performed as described previously [22]. Briefly, EnSC were seeded at a clonal density of 30 cells/cm^2^ on 1 mg/ml fibronectin-coated 6-well plates and cultured in 10% DMEM/F12 medium supplemented with basic fibroblast growth factor (bFGF; 10 ng/ml; Merck Millipore, Watford, UK) for 10 days with a 50 % media change on day 7. Colonies were monitored microscopically to ensure that they were derived from single cells. After 10 days, colonies were washed with PBS and fixed in 4% formaldehyde for 10 min at room temperature (RT) before staining with hematoxylin for 4 min. Colonies were visualized on EVOS FL AUTO microscope (Life Technologies, Paisley, UK). Clusters of ≥ 50 cells were counted and cloning efficiency (%) was calculated as the number of colonies formed/number of cells seeded ×100.

### Immunohistochemistry

Endometrial biopsies were fixed overnight in 10% neutral buffered formalin at 4°C and embedded in Surgipath Formula ‘R’ paraffin using the Shandon Excelsior ES Tissue processor (ThermoFisher). Tissues were sliced into 3 μM sections on a microtome and adhered to coverslips by overnight incubation at 60°C. Deparaffinization, antigen retrieval, antibody staining, hematoxylin counter stain and DAB color development were fully automated in a Leica BondMax autostainer (Leica BioSystems). Tissue sections were labelled for anillin (AMAB90660, Sigma) using a 1:100 dilution. Stained slides were de-hydrated, cleared and cover-slipped in a Tissue-Tek Prisma Automated Slide Stainer, model 6134 (Sakura Flinetek Inc. CA, USA) using DPX coverslip mountant. Bright-field images were obtained on a Mirax Midi slide scanner with a 20× objective lens, and 3 randomly selected areas of interest underlying the luminal epithelium were captured using Panoramic Viewer v1.15.3 (3DHISTECH Ltd, Budapest, Hungary). Each field was divided manually into 3 compartments: stroma, glandular epithelium, and luminal epithelium. ImageJ image analysis software (Rasband, W. S., ImageJ, National Institutes of Health) was used to quantify anillin^+^ cells with staining intensity manually determined by background thresholding. The percentage of anillin+ cells was calculated (i.e. anillin^+^ cells/total cells ×100) in each field of view and for each cellular compartment.

### Immunofluorescence microscopy

CFUs and EnSC monolayers in 6-well plates were fixed in 4% paraformaldehyde, permeabilized with 0.5% Triton X-100 (Sigma Aldrich) and blocked with 1% BSA/PBS. Cells were probed with mouse anti-anillin monoclonal primary antibody (1:100, Sigma-Aldrich) overnight at 4°C. Labelled cells were stained with Alexa-Fluor™ 594 anti-mouse secondary antibody (1:1000, Invitrogen) for 1 hour at RT and counterstained with VECTASHIELD antifade mounting medium with DAPI (Vector Laboratories, Peterborough, UK). Images were captured on an EVOS FL AUTO fluorescence microscope. Anillin^+^ cells were quantified in 3 randomly selected fields relative to total number of DAPI-stained cells using ImageJ image analysis software. Tissue sections were double labelled overnight at 4°C with antibodies against anillin (mouse or rabbit) and either CD34 (1:250, Abcam, Cambridge, UK), CD56 (1:200, Leica Biosystems), or CD163 (1:200, Abcam). Labelled cells were stained with Alexa Fluor™ 594 anti-mouse secondary antibody and Alexa Fluor™ 488 anti-rabbit secondary antibody. Tissue autofluorescence was eliminated using the TrueVIEW^®^ autofluorescence quenching kit (Vector Laboratories) according to manufacturer’s instructions. Tissue sections were stained with ProLong Gold antifade mounting medium with DAPI (Invitrogen) and examined using an EVOS FL Auto fluorescence microscope (Life Technologies).

### RNA-sequencing and analysis

Total RNA was extracted from CFUs and standard cultures using the AllPrep DNA/RNA Micro Kit (Qiagen, Manchester, UK) following the manufacturer’s protocol. RNA concentration was assessed using the Qubit RNA BR assay and RNA quality was analyzed on an Agilent 2100 Bioanalyzer (Agilent Technologies). Libraries were prepared using the TruSeq RNA Library preparation kit V2 (Illumina, Cambridge, UK) and sequenced on HiSeq 4000 (Illumina) with 75 bp paired-end reads at the Wellcome Trust Centre for Human Genetics (Oxford, UK). Transcriptomic maps of paired-end reads were generated using Bowtie-2.2.3, Samtools-0.1.19, and Tophat-2.0.12 against the hg19 reference transcriptome. Transcript counts were assessed by HTSeq-0.6.1 using the reverse strand setting and intersection non-empty mode and counts were assigned to Ensembl gene IDs. Differential gene expression was evaluated with the DESeq2 package v1.28.1 in R, using the Wald test and Benjamini and Hochberg FDR corrected *P* < 0.05. Principal component analysis on the log2-normalised count data was performed in MATLAB R2020b. RNA-seq library were analyzed using Euclidean distance were plotted with pheatmap package v1.0.12 in R. RNA-seq data were deposited in the Gene Expression Omnibus (http://www.ncbi.nlm.nih.gov/geo/), accession number GSE159266.

### Single-cell transcriptome analyses

Analysis of previously reported single-cell RNA-seq data of midluteal endometrial biopsies (GSE127918) was performed with Seurat R package v3.2.2. Single cells were assigned to phases of the cell cycle using the CellCycleScoring function. Marker genes were obtained with the FindMarkers function. Heatmaps of gene expression were generated with the gplots package v3.1.0. Ligand-receptor analyses were performed with CellphoneDB (https://www.CellPhoneDB.org.) [23]. Single-cell transcriptome data of the maternal-fetal interface, were obtained from ArrayExpress (E-MTAB-6701) and used for cell cycle scoring and differential gene expression analysis. Informative genes were visualized using an on-line browsing tool (https://maternal-fetal-interface.cellgeni.sanger.ac.uk/).

### Statistical Analysis

Data analyses were performed using the statistical package Graphpad Prism 6 and R v4.0.2. Mann-Whitney U test, student *t*-test, and one-way ANOVA on ranks (Kruskal-Wallis) test were used as appropriate. Spearman rank-order correlation coefficient was calculated to measure the strength and direction of association between two variables. *P* < 0.05 was considered statistically significant.

## Results

### Characterization of hPMC in midluteal endometrium

We recently reported on single-cell RNA-sequencing analysis of midluteal endometrial biopsies, i.e. coinciding with the window of implantation [8]. A notable observation was the presence of a discrete cell population comprising 1.8% of all non-immune stromal cells. This population was designated ‘highly proliferating mesenchymal cells’ (hPMC) based on 127 upregulated genes (FDR corrected *P* < 0.05) involved in cell cycle progression. In agreement, gene ontology analysis yielded terms such as ‘cell cycle’ (FDR corrected *P* = 6.3 × 10^-66^), ‘cell division’ (FDR corrected *P* = 4.77 × 10^-41^), and ‘regulation of mitotic cell cycle’ (FDR corrected *P* = 1.41 × 10^-22^). To explore the nature of hPMC, we first computationally determined the cell cycle status of all cells in midluteal endometrium (Fig 1A). The hPMC population consisted exclusively of cells in either S-phase (40%) or G2/M-phase (60%) (Fig. 1A). By contrast, the majority of EnSC and endometrial epithelial cells (EpC) were assigned to G1 (55% and 63%, respectively). The abundance of endothelial cells (EC) and immune cells (IC) in S or G2/M phase was 58% and 72%, respectively (Fig. 1A). Next, we compared gene expression in hPMC assigned to the G2/M phase to all other cells in G2/M across different populations. This analysis yielded 19 differentially expressed cell cycle genes (FDR corrected *P* < 0.05), all markedly upregulated in hPMC (Fig. 1B and Table S2). To determine the provenance of hPMC, we constructed a correlation matrix heatmap and dendrogram to show hierarchical clustering of different endometrial cell populations (Fig. 1C). The heatmap depicts the Euclidian distance in gene expression between cell populations whereas the distance between the dendrogram branches reflects clustering according to the complete linkage method. hPMC clustered with EC and EnSC, but not with EpC or IC, suggesting shared provenance (Fig. 1C).

**Figure 1.**
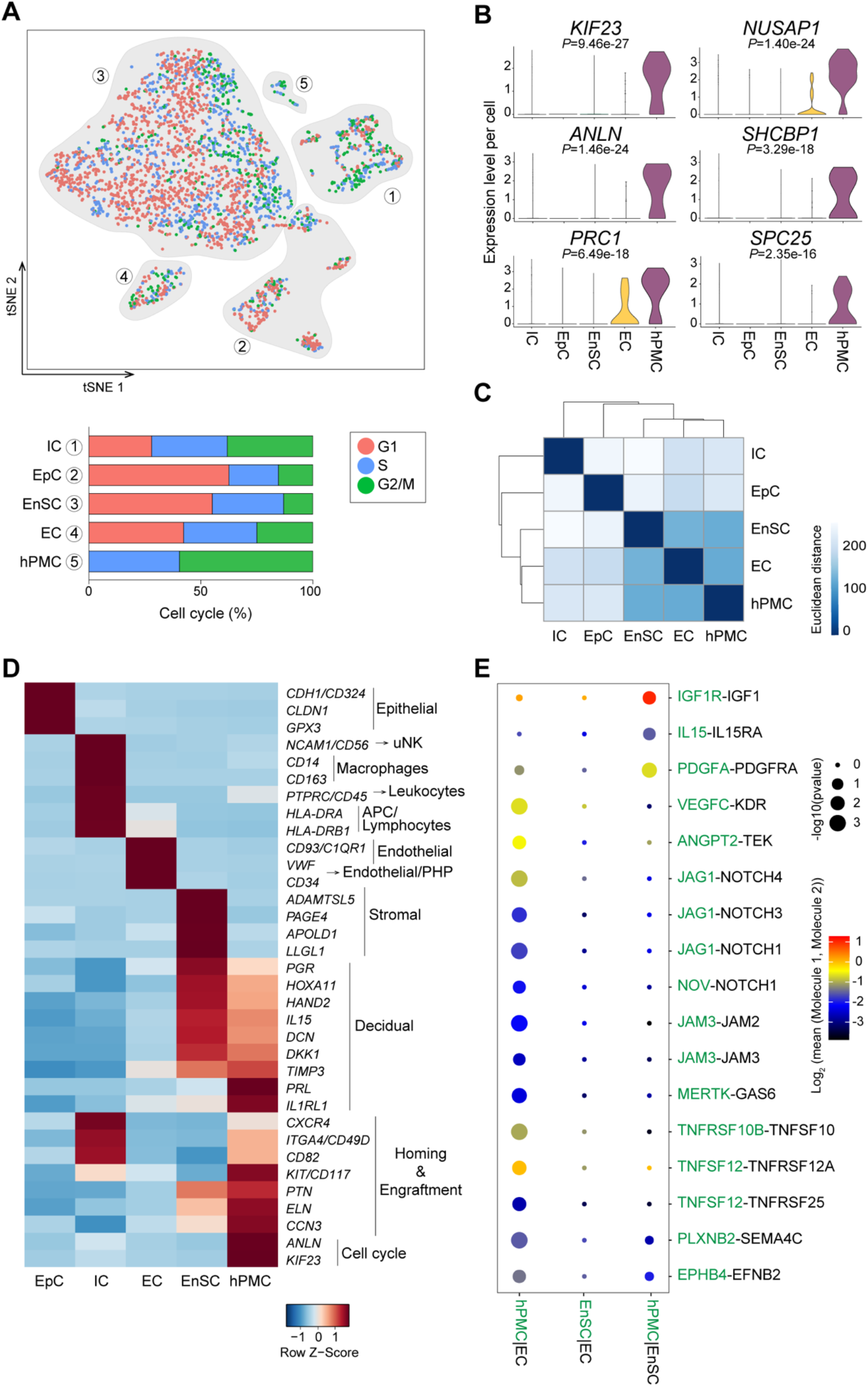
Transcriptomic analysis of hPMC. **(A)** Top panel shows *t*-SNE projection of mid-secretory endometrial cell populations [epithelial cells (EpC), immune cells (IC), endothelial cells (EC), stromal cells (EnSC) and highly proliferative mesenchymal cells (hPMC)] color-coded to indicate cells in different phases of the cell cycle (G1, S and G2/M). Lower panel shows the proportion of cells in the different phases of the cell cycle for each population. **(B)** Violin plots showing the relative expression in G2/M phase of 6 genes highly enriched in hPMC. FDR corrected *P*-values are also shown. **(C)** Heatmap showing Euclidean distance between different cell populations based on average expression values. **(D)** Heatmap showing relative expression (z-score) of genes used to characterize hPMC. Red and blue represent high and low gene expression, respectively, as indicated by the color key. **(E)** Overview of selected ligand–receptor interactions generated by Cellphone DB tool. FDR corrected *P*-values are reflected by circle sizes. The means of the average expression level of interacting molecule 1 in cluster 1 and interacting molecule 2 in cluster 2 are indicated by the color key.

Mining of gene expression data showed that hPMC do not express genes that encode for canonical IC, EC, or EpC markers (Fig. 1D). They also do not express several EnSC-specific genes that are constitutively expressed across the menstrual cycle (e.g. *ADAMTSL5*, *PAGE4*, *APOLD1* and *LLGL1*) (Fig. 1D and Fig. S1). However, hPMC share with EnSC the expression, of multiple decidual genes, including *DCN* (coding decorin), *DKK1* (dickkopf WNT signaling pathway inhibitor 1), *IL15* (interleukin 15), and *TIMP3* (TIMP metallopeptidase inhibitor 3) [8]. *PRL* (prolactin), a widely used marker gene of decidualizing EnSC *in vitro* [3], is conspicuously enriched in hPMC *in vivo*. Another notably enriched decidual gene is *IL1RL1*, which codes both the transmembrane and soluble IL-33 receptors (termed ST2L and sST2, respectively) [7]. hPMC also express key decidual transcription factors, including *PGR* (coding the progesterone receptor) [24], *HAND2* (heart and neural crest derivatives expressed 2) [25], and *HOXA11* (homeobox A11) [26], indicating commitment to decidual differentiation (Fig. 1D). On the other hand, hPMC share with IC genes involved in chemotaxis (e.g. *CXCR4* and *KIT*) and trans-endothelial migration (e.g. *ITGA4* and C*D82*) [27]. However, unlike IC, hPMC also express genes coding for secreted factors implicating in MSC self-renewal and engraftment (Fig. 1D), including tropoelastin (*ELN*) [28], pleiotrophin (*PTN*) [29], and cellular communication network factor 3 (*CCN3*) [30]. Next, we used CellPhoneDB to explore putative interactions between hPMC, EC and EnSC. This computational tool takes into account the subunit architecture of both ligands and receptors in heteromeric complexes [13,23]. As shown in Figure 1E and Table S3, CellPhoneDB predicted many more enriched receptor-ligand interactions between EC and hPMC than with EnSC. Specifically, hPMC selectively encode ligands that bind various EC receptors, including NOTCH 1, 3 and 4, TIE2 receptor (TEK), and VEGFR (KDR). On the other hand, activation of insulin-like growth factor 1 receptor (IGF1R) is predicted to play a role in chemotaxis of hPMC within the stromal compartment (Fig. 1E). Intriguingly, a recent study reported activation of IGF1R by CXCL14 [31], an orphan chemokine abundantly expressed in midluteal endometrium [32]. Taken together, the data suggest that hPMC are trans-endothelial migratory cells committed to decidual differentiation.

### Spatiotemporal distribution of hPMC in luteal phase endometrium

Cross-referencing of potential hPMC markers with the Human Protein Atlas (http://www.proteinatlas.org) revealed the presence of rare anillin^+^ cells scattered throughout the endometrium [33]. Anillin, encoded by *ANLN*, is an actin-binding protein involved in cytokinesis, cell growth and migration [34]. To examine if anillin could serve as a marker for hPMC, immunohistochemistry was performed on 61 LH-timed endometrial biopsies obtained across the luteal phase. This analysis confirmed that anillin^+^ cells are rare and dispersed throughout the stroma, although frequently enriched around the terminal spiral arterioles (Fig. 2A). Unexpectedly, some anillin^+^ cells were also found in glandular and luminal epithelium (Fig. 2A). The average abundance of anillin^+^ cells [median (interquartile range, IQR)] in the stromal compartment across the luteal phase was 0.6 (1.3)% versus 0.4 (0.7)% and 0.2 (0.6)% in the luminal and glandular epithelium, respectively. In contrast to glandular or luminal epithelium, anillin^+^ cells increased significantly in the stroma upon progression from early to late-secretory phase of the cycle (Fig. 2B). Quantification of anillin^+^ cells showed a strong correlation between the three main cellular compartments (Fig. 2C), especially between the stroma versus glandular epithelium (*r* = 0.7, *P* < 0.0001, Spearman’s rank test) and glandular versus luminal epithelium (*r* = 0.7, *P* < 0.0001). Nevertheless, proportionally many more anillin^+^ cells reside in the stroma when compared to glandular or luminal epithelium (Fig. S2).

**Figure 2.**
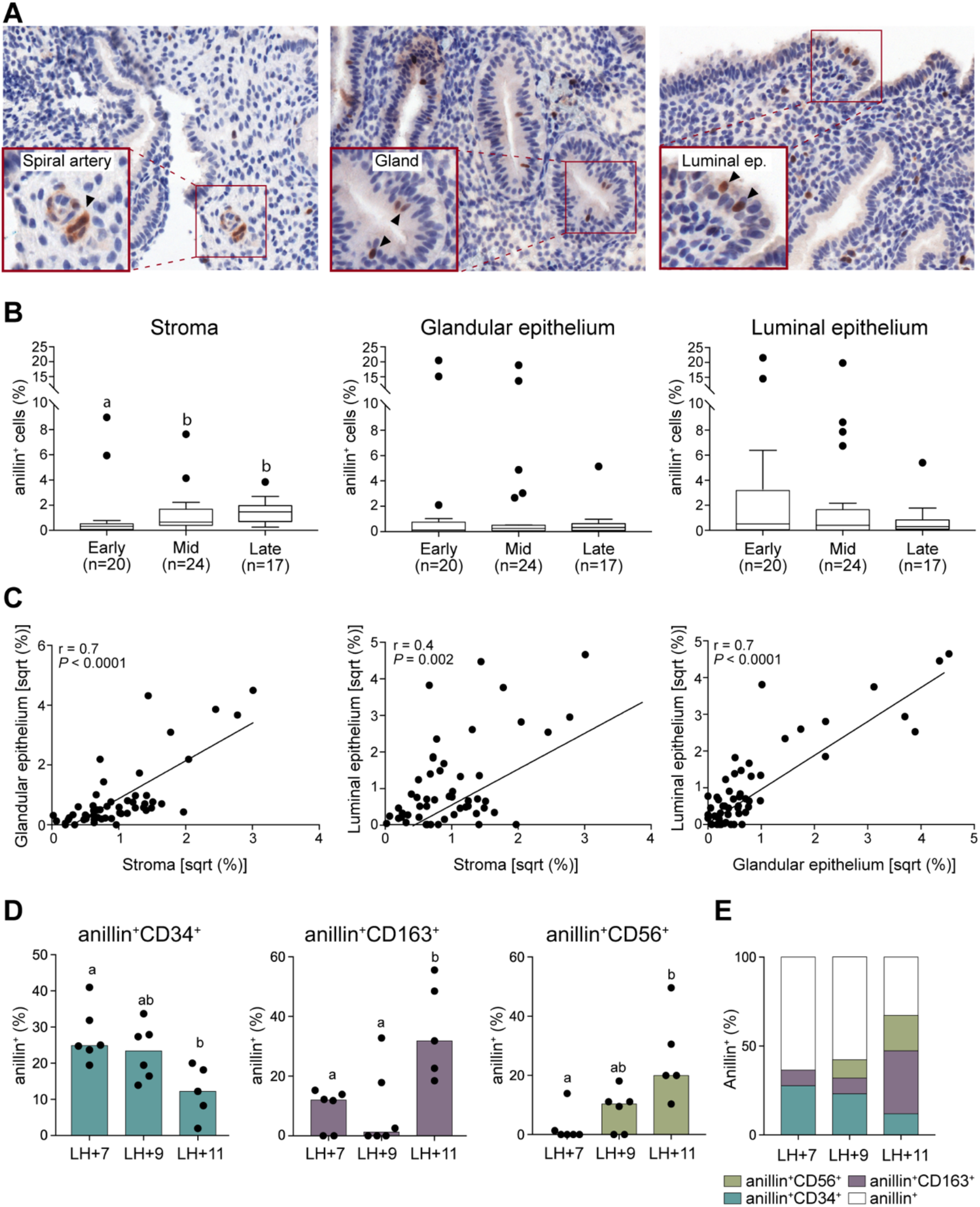
Spatiotemporal distribution of anillin^+^ cells in peri-implantation endometrium. **(A)** Examples of anillin^+^ cells in endometrial stroma and glandular and luminal epithelium. **(B)** Quantification of anillin^+^ cells in different endometrial compartments during the early-(LH+5/6), mid-(LH+7/9) and late-secretory (LH+10/11) phase of the cycle. The number of biopsies analyzed at each timepoint is indicated. Different letters above box plots indicate that groups are significantly different from each other at *P* < 0.05 (one-way ANOVA on ranks test). **(C)** Spearman’s rank correlation of anillin^+^ cells in endometrial stroma, glandular epithelium and luminal epithelium. **(D)** Quantification of anillin^+^CD34^+^ hematopoietic/endothelial precursor cells, anillin^+^CD163^+^ macrophages, and anillin^+^CD56^+^ uNK cells in peri-implantation endometrium. Different letters above the box plots indicate that groups are significantly different from each other at *P* < 0.05 (one-way ANOVA on ranks test). **(E)** Relative proportions of anillin^+^ IC and hPMC (non-immune anillin^+^ cells) during the midluteal window of implantation (LH+7/9) and late-secretory phase (LH+11).

We reasoned that the time-dependent increase in anillin^+^ cells in the stroma could be accounted for by accumulation of proliferating uNK cells or macrophages prior to menstrual shedding [15]. To test this conjecture, 21 endometrial biopsies were subjected to double-labelling immunofluorescence microscopy to quantify co-expression of anillin with CD163 or CD56, macrophage and uNK cell markers, respectively [6,35]. To assess the contribution of proliferating leukocyte and EC precursors, the abundance of anillin^+^ cells co-expressing CD34 was also quantified (Fig. S3) [36]. The samples were selected to coincide with the implantation window (LH+7 and LH+9) or the late-secretory phase of the cycle (LH+11). Anillin^+^ CD56^+^ cells were noticeably absent in all but one sample at LH+7 but their abundance increased markedly upon transition to the late-secretory phase in parallel with anillin^+^CD163^+^ macrophages (Fig. 2D). The reverse pattern was observed for anillin^+^CD34^+^ cells. At the start of the implantation window (LH+7), 61% of anillin^+^ cells did not express IC markers, but this dropped to 32% by the late-secretory phase of the cycle (Fig. 2E).

### hPMC confer endometrial clonogenicity

To determine if hPMC confer endometrial clonogenicity, we first established paired colony forming unit (CFU) assays and standard primary EnSC cultures from 3 biopsies (Fig. 3A). After 7 days, the abundance of anillin^+^ cells [median (IQR)] in standard cultures was 1.7 (1.9)% versus 17 (7.7)% in clonal assays (*P* < 0.05, Mann Whitney U test). To test whether culturing leads to loss of proliferative capacity, anillin^+^ cells were quantified using immunofluorescence microscopy in CFU assays after 3, 6 or 9 days in culture (Fig. 3B). Notably, the abundance of anillin^+^ clonal cells on day 3 was 68 (17.7)% but this dropped significantly by day 9 to 15.1 (9.6)% (*P* < 0.0002, Kruskal-Wallis test). Next, we examined if the abundance of anillin^+^ cells *in vivo* is a measure of endometrial clonogenicity. CFU assays were established from 47 midluteal endometrial biopsies and, in parallel, anillin^+^ cells were quantified in the stromal compartment using immunohistochemistry (Fig. 3C). A significant correlation was observed between the percentage of anillin^+^ stromal cells *in vivo* and CFU activity of freshly isolated EnSC *in vitro* (*r* = 0.49, *P* < 0.0004, Spearman’s rank test).

**Figure 3.**
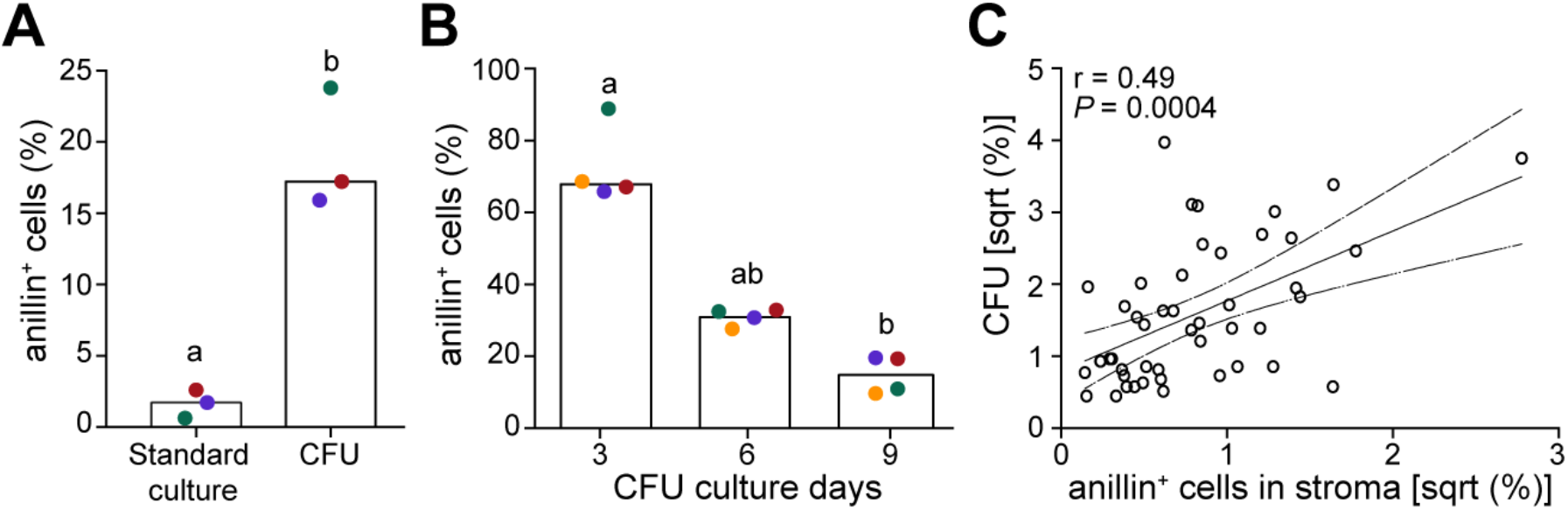
hPMC confer clonogenicity to midluteal superficial endometrium. **(A)** Percentage of anillin^+^ cells in primary EnSC cultures and CFU assays. Bar graphs show median of 3 biological repeat experiments. **(B)** Percentage of anillin^+^ cells in CFU assays on day 3, 6 and 9 of culture. Bar graphs show median of 4 biological repeat experiments. Different letters above the box plots indicate that groups are significantly different from each other at *P* < 0.05 (one-way ANOVA on ranks test). **(C)** Spearman’s rank correlation of anillin^+^ cells in the endometrial stromal compartment *in vivo* and CFU activity of freshly isolated EnSC in 47 paired midluteal endometrial biopsies.

### Transcriptomic profiling of cultured clonogenic and resident stromal cells

Previous studies reported that CFU assays are more efficient when established from SUSD2^+^ perivascular cell (PVC) fraction when compared to non-perivascular (SUSD2^-^) EnSC [22,37]. We reasoned that if hPMC are derived from circulating progenitor cells and account for endometrial clonicity during the implantation window, the transcriptomic profiles of clonogenic cells in culture should exhibit less variability when compared to cultured resident PVC or EnSC that have been exposed to iterative menstrual cycles *in vivo*. To test this hypothesis, we used magnetic-activated cell sorting to isolate SUSD2^+^ PVC and SUSD2^-^ EnSC from 3 different biopsies and subjected both cell fractions to standard cultures as well as CFU assays (Fig. 4A). For clarity, clonogenic cells established from PVC were termed endometrial MSC (eMSC) whereas the term ‘transit amplifying’ (TA) cells was coined to clonal cells in the EnSC fraction. After 10 days in culture, total RNA was extracted and subjected to RNA-seq analysis. Principal component (PC) analysis separated standard EnSC and PVC cultures from eMSC and TA in, PC1 (Fig. 4B). Notably, all six libraries from clonal cells clustered tightly together. Although PVC and EnSC also clustered together, there was considerable divergence in their transcriptional profiles between samples (Fig. 4B). Thus, in keeping with our hypothesis, cultured resident cells exhibit much greater interpatient transcriptional variability when compared to clonogenic cells.

**Figure 4.**
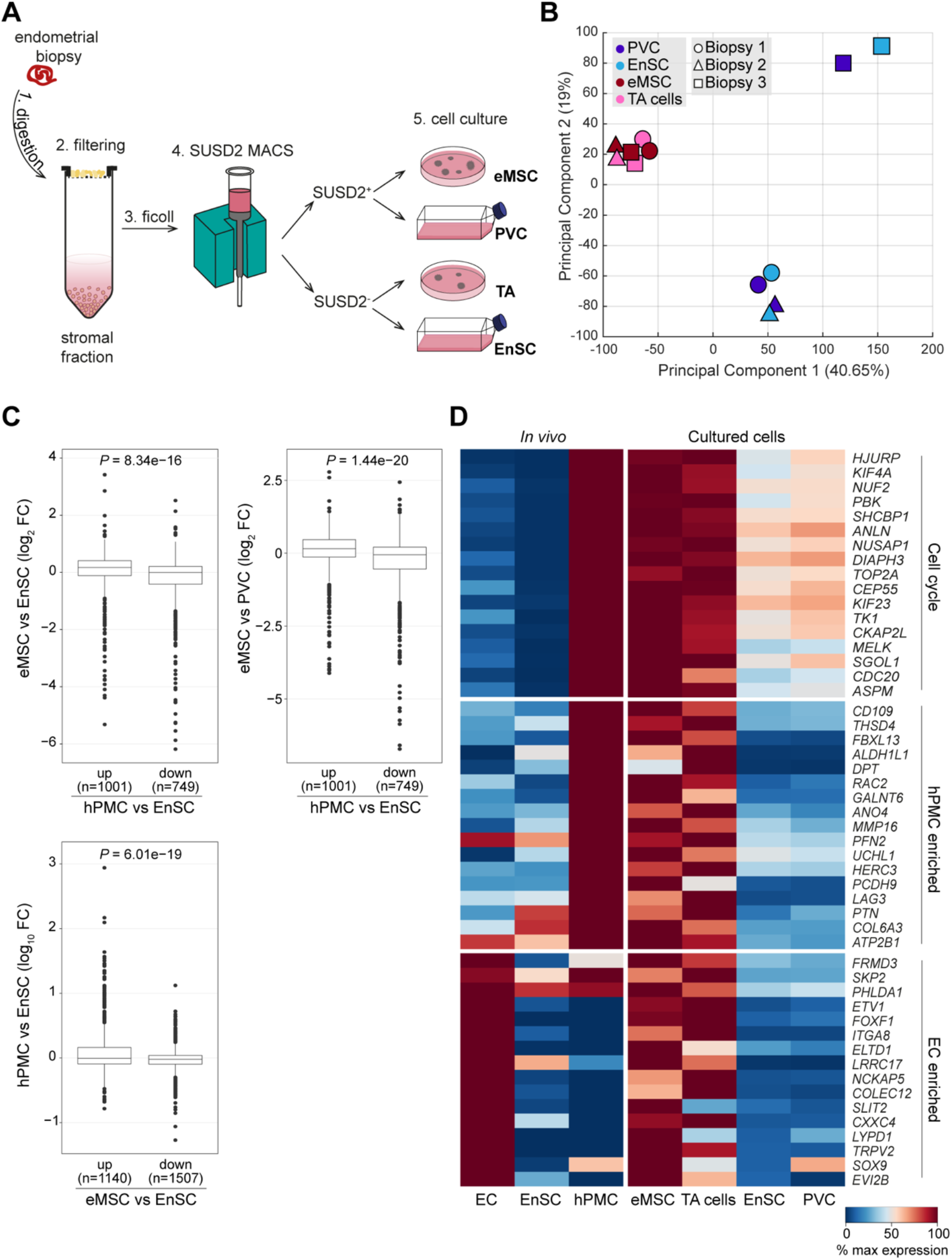
A shared gene signature between clonogenic cells in culture and hPMC *in vivo*. **(A)** Schematic depiction of the experimental design. After 10 days in culture, total RNA from 3 biological repeat experiments was subjected to bulk RNA-sequencing. **(B)** Principal-component analysis of RNA-seq data of cultured clonogenic cells (eMSC/TA) and resident cells (EnSC/PVC). **(C)** Upper panels: box plots showing relative expression in eMSC versus EnSC or eMSC versus PVC of genes either enriched or depleted in hPMC when compared to EnSC. Lower panel: box plot showing relative expression in hPMC versus EnSC of genes either significantly enriched or depleted in eMSC when compared to EnSC and PVC. For *in vitro* bulk RNA-seq data, DEGs were defined as = 2 fold-change (FC) and FDR corrected *P* < 0.05. For *in vivo* scRNA-seq data, DEGs were defined as *P* < 0.05 and expression in = 10% of cells. **(D)** Composite heatmap showing relative expression of indicated genes in EC, EnSC and hPMC *in vivo* and eMSC, TA cells, EnSC and PVC *in vitro*. The color key indicates the level of gene expression relative to the highest expressing cell population *in vivo* or *in vitro*.

Based on > 2 fold-change in gene expression and FDR corrected *P* < 0.05, only two transcripts (*STMN3* and *ITGBL1)* were differentially expressed between eMSC and TA cells, indicating shared origins. By contrast, 3,460 genes were differentially expressed between clonogenic cells (eMSC plus TA cells) and resident stromal cells (PVC plus EnSC). Out of the 3,460 genes, 81.8% were down-regulated in clonal cells. The heatmap in Figure S4A depicts the 10 most highly enriched or depleted genes in clonal cells when compared to EnSC and PVC. In keeping with the results of a previous study [37], a total of 216 differentially expressed genes (DEG) were identified between cultured PVC and EnSC. Interesting, some genes enriched in PVC versus EnSC appeared also higher expressed in eMSC and, to a lesser extent, in TA cells, although statistical significance was often not reached (Fig. S4A). An UpSet plot was used to depict the intersections of DEG in different comparisons (Fig. S4B).

Clonogenic cells rapidly undergo spontaneous differentiation and senescence in culture [38]. Nevertheless, differences in gene expression between hPMC versus EnSC *in vivo* were partially maintained in eMSC when compared to EnSC (*P* = 8.3 × 10^-16^, *t*-test) or PVC (*P* = 1.4 10^-22^) *in vitro* (Fig. 4C and Table S4). Conversely, differences in gene expression between clonal and resident cells *in vitro* were also partially maintained between hPMC and EnSC *in vivo* (*P* = 6.1 × 10^-19^) (Fig. 4C). Mining of the data provided further evidence that hPMC are the progenitors of colony-forming cells in culture. As anticipated, proliferation-associated genes were higher expressed in eMSC/TA cells when compared to cultured EnSC or PVC (Fig. 4D), including multiple genes implicated in cancer stem cells (e.g. *PBK*, *NUSAP1*, *MELK*, *CDC20* and *ASPM*) [39–42]. Further, several MSC-related genes enriched in hPMC were also selectively enriched in eMSC/TA cells. For example, *ALDH1L1*, encoding aldehyde, dehydrogenase 1 family member L1, and *CD109* are putative biomarkers of cancer stem-like cells [43,44], whereas profilin 2 (*PFN2*) promotes migration, invasion and stemness of HT29 human colorectal cancer stem cells [45]. Matrix metallopeptidase 16 (*MMP16*) controls the migration of human cardiomyocyte progenitor cells [46]. Notably, eMSC, and to a lesser extent TA cells, also express an EC-like gene signature, albeit devoid of canonical endothelial marker genes (Fig. 4D). Multiple genes in this signature, such as *SKP2*, *ITGA8*, *SLIT2* and *SOX9*, have been shown to be implicated in different stem/progenitor cell niches [47–50].

### Loss of hPMC in recurrent pregnancy loss

We reported previously that recurrent pregnancy loss, defined here as 3 or more miscarriages, is associated with loss of eMSC and TA cells, as assessed by CFU assays on freshly isolated PVC and EnSC, respectively, from midluteal biopsies [14]. Hence, we examined if this prevalent reproductive disorder is also associated with loss of anillin^+^ hPMC *in vivo*. To mitigate against interference from anillin^+^ IC, the analysis was restricted to endometrial biopsies obtained on LH+7, a timepoint when anillin^+^ CD56^+^ cells are virtually absent (Fig. 2D). A total of 30 biopsies from RPL patients (n = 15) and control subjects (n = 15) were processed for immunofluorescence microscopy to quantify anillin^+^ cells in the stromal compartment as well anillin^+^ cells co-expressing CD163 or CD34. While the percentage of anillin^+^CD163^+^ cells did not differ between the two clinical groups (Fig. 5A), a significant increase in anillin^+^CD34^+^ cells was observed in RPL patients when compared to control subjects (*P* < 0.05, Mann-Whitney test; Fig. 5B). Conversely, RPL was associated with a reciprocal reduction in hPMC, i.e. anillin^+^ cells that do not express IC markers (*P* < 0.05, Mann-Whitney U test; Fig. 5C). The abundance of hPMC population was noticeably more variable in tissue samples from RPL patients when compared to control subjects (Fig. 5C). Further, hPMC made up > 50% of all anillin^+^ cells in 14 out of 15 control samples compared to only 7 out of 15 samples in the RPL group (*P* = 0.01, Fisher’s exact test).

**Figure 5.**
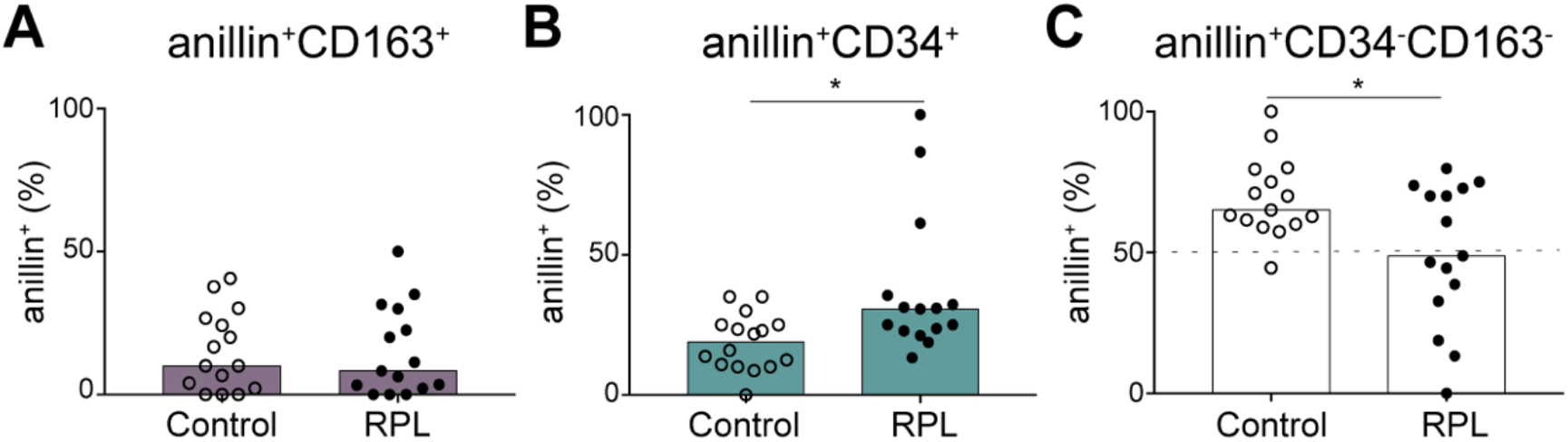
Loss of hPMC in recurrent pregnancy loss (RPL). The percentage of anillin^+^ cells co-expressing **(A)** CD163 (macrophages) or **(B)** CD34 (hematopoietic / endothelial precursor cells) was quantified by immunofluorescent microscopy in endometrial biopsies obtained on LH+7 from control subjects (n=15) and RPL patients (n=15). **(C)** The relative abundance of hPMC (i.e. anillin^+^ cells not co-expressing CD34 or CD163) in endometrial biopsies from control and RPL subjects. * indicates *P* < 0.05 (Mann-Whitney U test).

### hPMC map to a distinct decidual subpopulation in pregnancy

Next, we set out to determine the potential fate of hPMC in early pregnancy. To do this, we made use of the recently published single-cell atlas of the maternal-fetal interface in early gestation [13]. Anatomically, the maternal decidua consists of two distinct layers, the decidua compacta near the surface epithelium and the underlying decidua spongiosa, which contains hypersecretory glands. The two decidual layers contain 3 decidual subpopulations, designated dS1-3 (Fig. S5A) [13]. The decidua spongiosa harbors dS1 cells expressing *ACTA2* whereas the dS2 and dS3 cells in the decidua compacta share the expression of *DCN* (Fig. S5B). Notably, dS3 cells constitute a minor subpopulation, representing only 7.2% of all decidual stromal cells at the maternal-fetal interface. Computational analysis indicated that 7% of dS3 cells are in S or G2/M phase of the cell cycle compared to 2.6% and 3.1% of dS2 and dS1 cells, respectively. Although dS3 cells are much less proliferative than hPMC, they nevertheless share a conspicuous gene signature (Fig. 6A and Table S5). This signature is not only enriched in several canonical decidual genes, including *PRL*, *IL1RL1*, and *CD82* (also known as *KAI1*) [7,24,51], but also in genes involved in glycolysis and mitochondrial function (Fig. 6B and Fig. S5B). Conversely, hPMC in midluteal endometrium and dS3 cells at the maternal-fetal interface in pregnancy are relatively depleted in genes implicated in cellular stress responses (Fig. 6B), including members of the AP-1 family of transcription factors (*FOS*, *FOSB*, *JUN*, and *JUNB*) and CEBPβ (*CEBPB*), which effects free radical signaling in decidual cells [52]. Based on this shared metabolic and decidual gene signature, we conclude that hPMC likely give rise in pregnancy to a distinct decidual subpopulation that engages with invading placental trophoblast in the decidua compacta.

**Figure 6.**
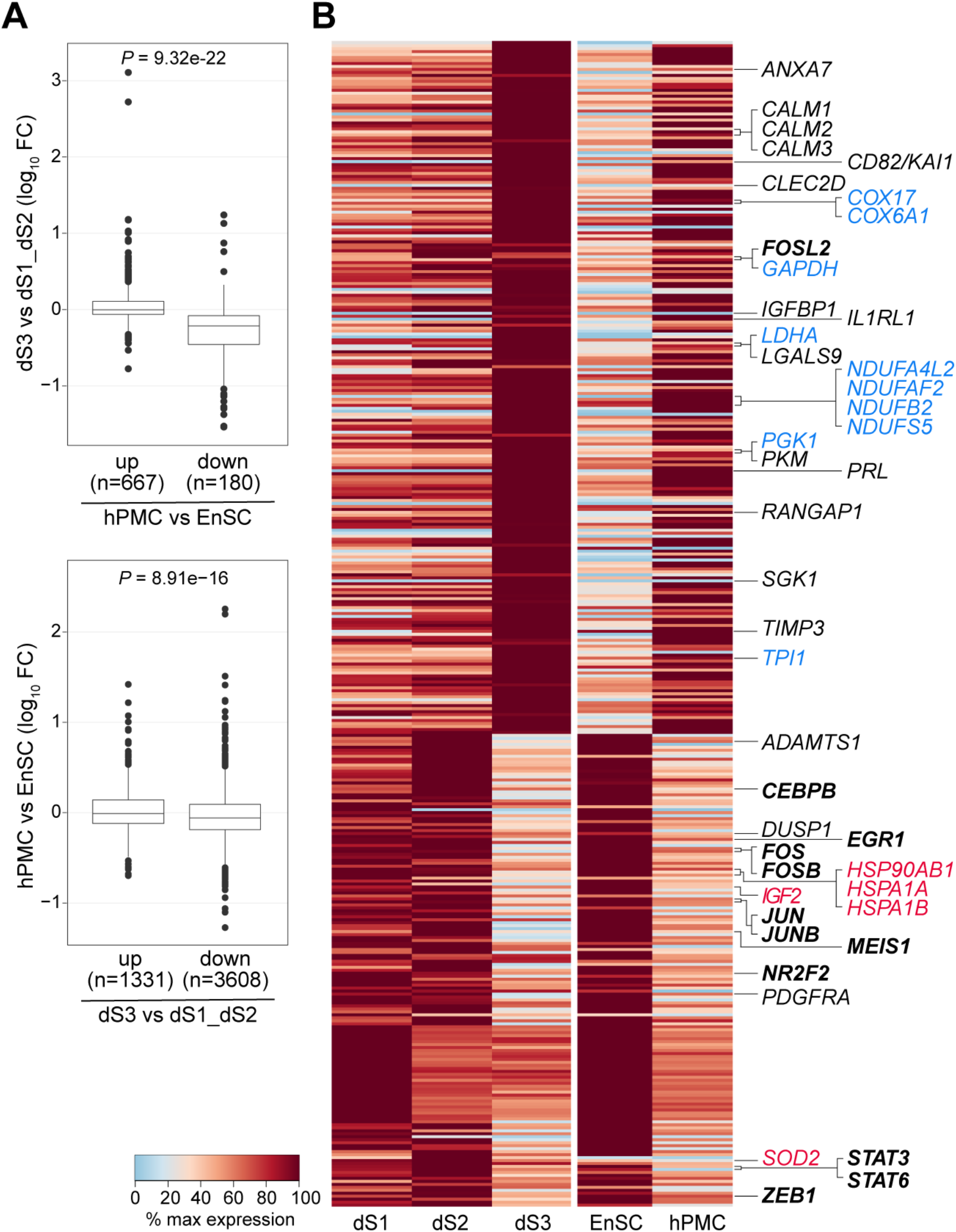
hPMC are putative precursors of a distinct decidual subpopulation in pregnancy. **(A)** Upper panel: box plots showing the relative expression in dS3 versus dS1 and dS2 subpopulations of genes either significantly (*P* < 0.05 and expression in > 10% of cells.) enriched or depleted in hPMC when compared to EnSC and EC in midluteal endometrial. Lower panel: box plots showing the relative expression in hPMC versus EnSC of genes either significantly enriched or depleted in dS3 cells when compared to dS1 and dS2 populations at the maternal-fetal interface in early pregnancy. (**B)** Composite heatmap showing relative expression of indicated genes in the 3 decidual subsets (dS1-3) at the maternal-fetal interface in early gestation and in EnSC and hPMC during the window of implantation. Annotated genes were color-coded to indicate stress-related genes (red), mitochondrial and metabolic genes (blue), or genes coding for known decidual factors (black) or transcription factors (bold black). The color key indicates the level of gene expression relative to the highest expressing cell population either in pregnancy (dS1-3) or during the implantation window prior to conception (EnSC and hPMC).

## Discussion

Decidual transformation of human endometrium upon embryo implantation requires differentiation of resident EnSC, immune clearance of acute senescent decidual cells, and rapid tissue growth. These processes occur in concert with coordinated trophoblast invasion [53]. Considering the magnitude of tissue remodeling required for pregnancy, poised progenitor and highly proliferative decidual precursor cells are likely critical for the formation of a robust maternal-fetal interface, as is the case in mice [20]. Here, we provide evidence of recruitment, engraftment and differentiation of hPMC/decidual precursor cells during the window of implantation in human endometrium. Transcriptional profiling highlighted the transitional nature of hPMC, exemplified on the one hand by expression of genes involved in chemotaxis, EC interactions, migration, and engraftment and, on the other, by expression of decidual transcription factors and marker genes. Human BDMSC are capable of differentiating into prolactin-producing decidual cells *in vitro* [54], further supporting our assertion that hPMC are equivalent to bone marrow-derived decidual precursor cells in pregnant mice. We demonstrate that hPMC share a conspicuous gene signature with dS3 cells identified recently in the decidua compacta of the maternal-fetal interface [13]. This common gene signature is defined by the expression of specific decidual marker genes, including *PRL*, *IL1RL1*, and *CD82* [7,24,51], enhanced expression of mitochondrial and metabolic genes, and the relative lack of stress-related genes when compared to either resident EnSC in midluteal endometrium or other decidual subsets present at the maternal-fetal interface in pregnancy. Further, a hallmark of dS3 cells in pregnancy is the expression of *HSD11B1* [13], which converts inactive cortisone into cortisol, a potent anti-inflammatory glucocorticoid [55]. Although our computational analysis showed that the proliferative capacity of dS3 cells is higher than that of other decidual subsets, it is markedly lower when compared to hPMC. It is plausible that recruitment of hPMC to the maternal-fetal interface diminishes rapidly upon invasion and plugging of the maternal terminal arterioles by extravillous trophoblast and subsequent physiological transformation of the spiral arteries [56–58]. If correct, the level of cellular plasticity in the decidual basalis, and thus protection against decidual senescence throughout gestation, may be determined during a relatively narrow window in early gestation.

Studies on the role of BDMSC in endometrial physiology have largely focussed on tissue regeneration [59–61]. Menstrual ‘injury’ and rising estradiol levels upregulate CXCL12, a potent chemotactic factor that mediates mobilization and homing of BDMSC by activating CXCR4 [62]. It is unclear whether the CXCL12/CXCR4 axis also mediates recruitment of BDMSC during the progesterone-dominant luteal phase. Notably, the expression of the evolutionarily related chemokine CXCL14 is at least a magnitude higher during the window of implantation when compared to CXCL12 [4,8]. CXCL14 is an orphan chemokine acting on uNK cells and immature dendritic cells but not T cells [63]. It also stimulates fibroblast proliferation and migration [64], as well as embryonic stem cell renewal [31]. The peri-implantation endometrium is further characterized by increased vascular permeability [65], which arguably facilitates extravasation of circulating hematopoietic and nonhematopoietic cells. BDMSC also give rise to EC progenitor cells in both human and murine endometrium [20,61,66,67]. This shared ontogeny with hPMC may account for our observation that clonogenic cells in culture spontaneously express EC-related genes.

Our characterization of bone marrow-derived decidual precursors in human endometrium has potentially far-reaching clinical implications. As mentioned, bone marrow transplantation experiments in mice provided elegant proof-of-concept evidence of the therapeutic potential of BDMSC, demonstrating that wild-type BDMSC restore uterine defects, enhance decidualization, and prevent pregnancy loss in Hoxa10^+/−^ mice [20]. We reported previously that recurrent pregnancy loss is associated with loss of endometrial clonogenicity in midluteal endometrium, measured by CFU assays on freshly isolated EnSC [14], and increased frequency of cycles with a pro-senescent decidual response [8]. We now demonstrate that a lack of decidual precursor cells accounts for the loss of endometrial clonogenicity in recurrent miscarriage. A history of miscarriage also increases the risk of preterm birth [68,69], the leading cause of neonatal death and morbidity worldwide [70]. Interestingly, while recurrent pregnancy, loss is associated with an overt pro-senescent decidual response [8,16], spontaneous preterm labor is linked to accelerated decidual ageing and loss of plasticity at the maternal-fetal interface [71,72]. Hence, it is plausible that treatments aimed at enhancing recruitment, engraftment and differentiation of hPMC in the endometrium prior to, or soon after, conception will not only reduce the risk of miscarriage but also prevent preterm labor in at risk women. In this context, a recent pilot trial demonstrated that sitagliptin, a dipeptidyl-peptidase IV inhibitor used in the management of diabetes, increases endometrial clonogenicity and decreases decidual senescence during the implantation window in recurrent miscarriage patients [73]. The efficacy of sitagliptin, and other potential interventions that target BDMSC and decidual precursor cells [27], in mitigating the risk of adverse pregnancy outcome warrants further evaluation in clinical trials.

## Conclusions

This study describes for the first time the presence of decidual precursor cells during the window of implantation in human endometrium. Based on our findings, we posit that decidual precursors are derived from circulating BDMSC, primed for exponential growth in early pregnancy, and integral to the formation of the decidua compacta of the maternal-fetal interface. Conversely, lack of decidual precursors in a conception cycle may trigger a concatenation of events, resulting in breakdown of the decidua in early gestation and miscarriage or, plausibly, accelerated ageing of the maternal-fetal interface and spontaneous preterm labor.

## Acknowledgments

This work was supported by funds from the Tommy’s National Miscarriage Research Centre and Wellcome Trust Investigator Award to J.J.B (212233/Z/18/Z).

We are grateful to all the women who participated in this research and those who facilitated their participation. We also thank the High-Throughput Genomics Group at the Wellcome Trust Centre for Human Genetics for the generation of sequencing data.

## Disclosure of Potential Conflicts of Interest

The authors indicate no potential conflicts of interest

## Notes

### Competing Interest Statement

The authors have declared no competing interest.

