## Supplementary Information for "Characterization of highly proliferative decidual precursor cells during the window of implantation in human endometrium"

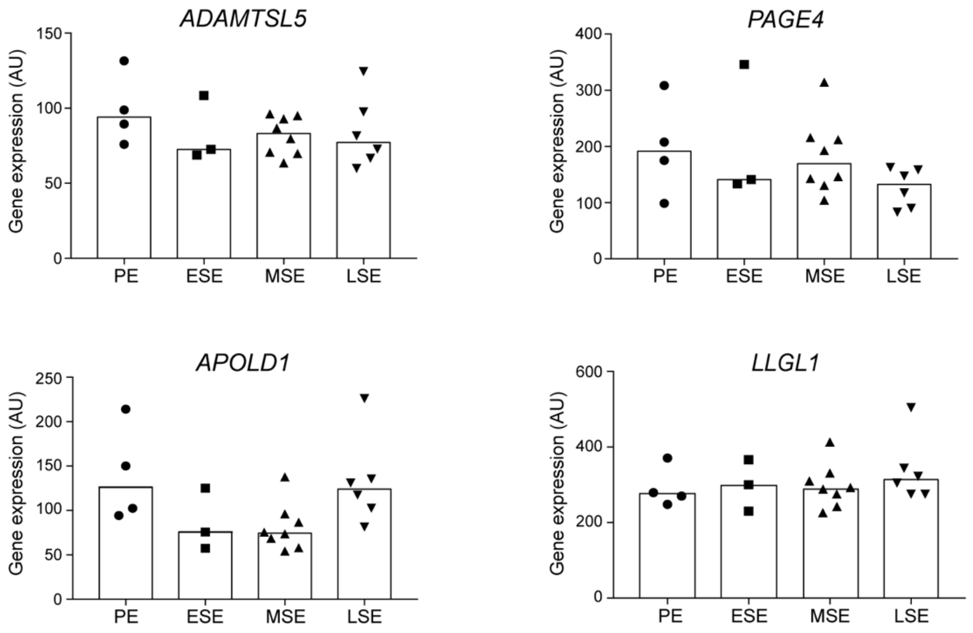


**Figure S1.** Cycle-independent EnSC-specific genes. EnSC marker genes were curated from midluteal endometrial scRNA-seq data (GSE127918) and their expression across the menstrual cycle examined by interrogating a publicly available microarray data set (GSE4888). Proliferative endometrium (PE, n=4); early secretory endometrium (ESE, n=3); mid-secretory endometrium (MSE, n=8) and late secretory endometrium (LSE, n=6).

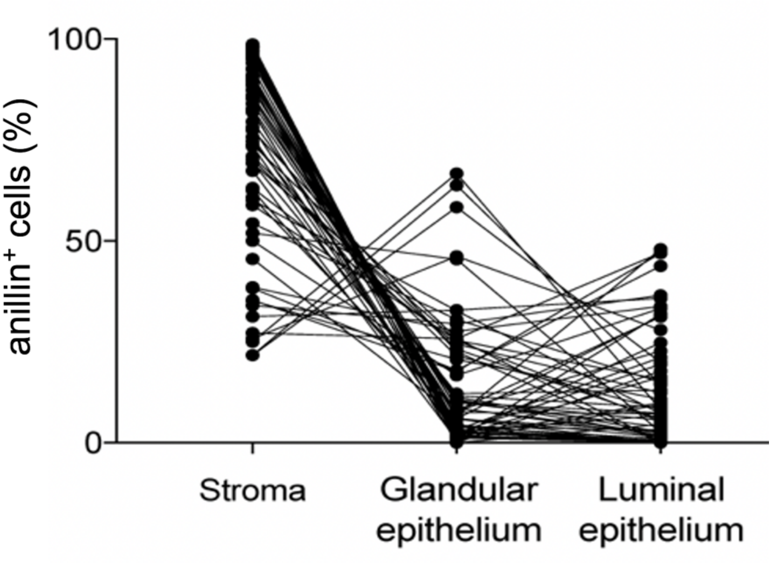


**Figure S2.** Relative distribution of anillin$^{+}$ cell in different endometrial compartments in 61 luteal phase endometrial biopsies

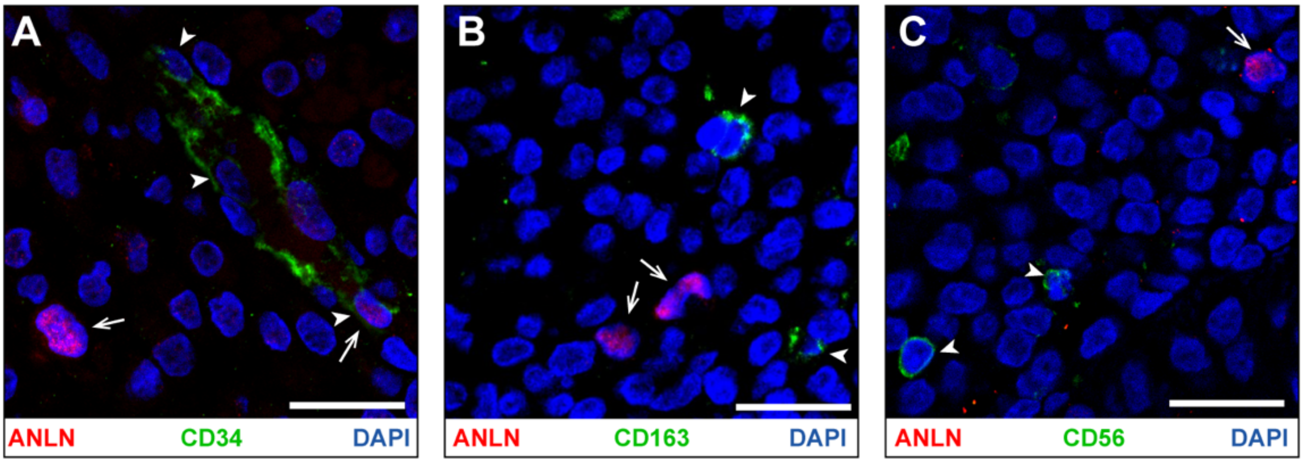


**Figure S3.** Examples of anillin$^{+}$ cells not expressing immune cell markers. Paraffin-embedded endometrial tissue sections were subjected to double immunofluorescence staining for anillin (ANLN) and CD34 (**A**), anillin and CD163 (**B**), and anillin and CD56 (**C**). Arrows indicate anillin$^{+}$ cells. Arrow heads indicate CD34$^{+}$, CD163$^{+}$ or CD56$^{+}$ cells. Scale bar = 20 μm.

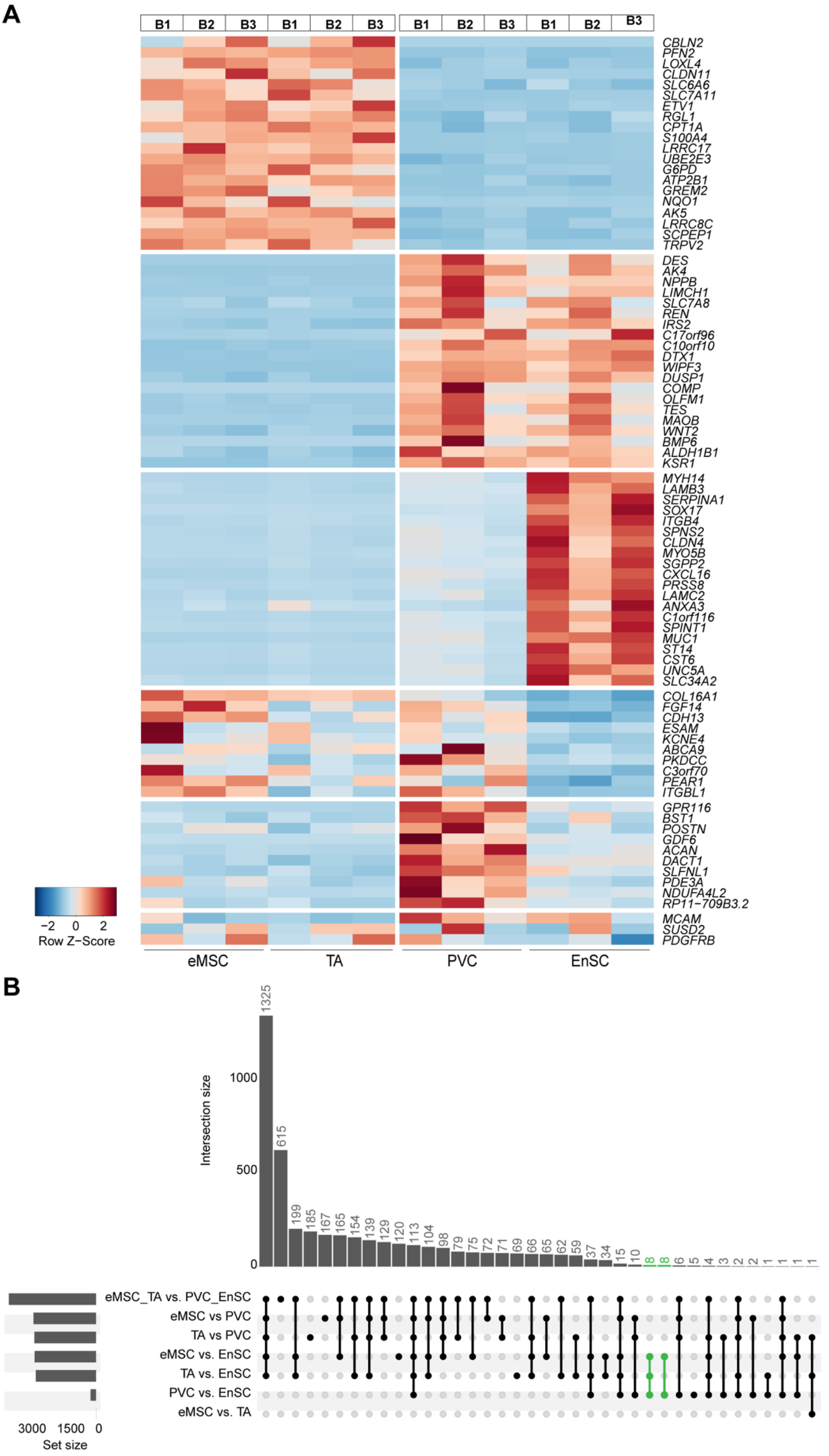
A
B1
B2
B3
CBLN2
PFN2
LOXL4
CLDN11
SLC6A6
SLC7A11
ETV1
RGL1
CPT1A
S100A4
LRRC17
UBE2E3
G6PD
ATP2B1
GREM2
NQO1
AK5
LRRC8C
SCPEP1
TRPV2
DES
AK4
NPPB
LIMCH1
SLC7A8
REN
IRS2
C17orf96
C10orf10
DTX1
WIPF3
DUSP1
COMP
OLFM1
TES
MAOB
WNT2
BMP6
ALDH1B1
KSR1
MYH14
LAMB3
SERPINA1
SOX17
ITGB4
SPNS2
CLDN4
MYO5B
SGPP2
CXCL16
PRSS8
LAMC2
ANXA3
C1orf116
SPINT1
MUC1
ST14
CST6
UNC5A
SLC34A2
COL16A1
FGF14
CDH13
ESAM
KCNE4
ABCA9
PKDCC
C3orf70
PEAR1
ITGBL1
GPR116
BST1
POSTN
GDF6
ACAN
DACT1
SLFNL1
PDE3A
NDUFA4L2
RP11-709B3.2
MCAM
SUSD2
PDGFRB
-2 0 2
Row Z-Score
eMSC
TA
PVC
EnSC
B
Intersection size
0
500
1000
1325
615
199
185
167
165
154
139
129
120
113
104
98
79
75
72
71
69
66
65
62
59
37
34
15
10
8
8
6
5
4
3
2
2
1
1
1
1
eMSC_TA vs. PVC_EnSC
eMSC vs PVC
TA vs PVC
eMSC vs. EnSC
TA vs. EnSC
PVC vs. EnSC
eMSC vs. TA
3000 1500 0
Set size

**Figure S4.** (**A**) Heatmap showing relative expression (z-score) of markers of cultured clonal endometrial cell populations (eMSC and TA cells) and resident subpopulations (PVC and EnSC). The relative expression of purported eMSC marker genes [i.e., *MCAM* (CD146), PDGFRB (CD140B), and SUSD2] are also shown (bottom of heatmap). The cultures were established from 3 independent biopsies, labelled B1-3. Red and blue represent high or low expression of a given marker gene, respectively, as indicated by the color key. (**B**) UpSet plot of intersections between sets of differentially expressed genes (FDR < 0.05, FC > 2) of eMSC and TA compared to each stromal subpopulation. The bar chart on the left indicates the total number of DEGs for each comparison. The upper bars indicates the intersection size between sets of DEGs. The matrix of solid and empty circles at the bottom illustrates indicate the cell populations for each intersection.

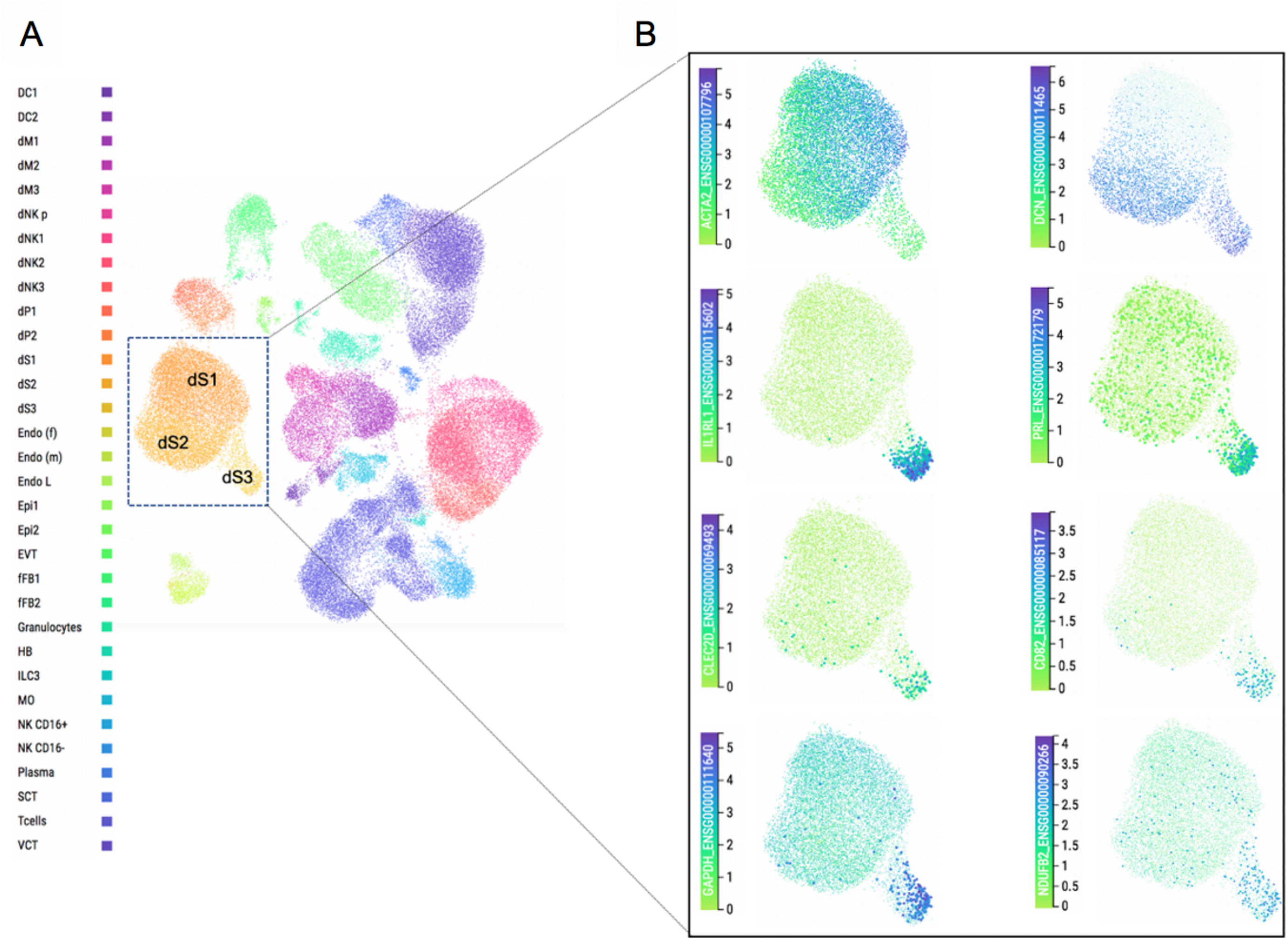


**Figure S5.** A shared gene signature between hPMC during the window of implantation and the decidual subpopulation dS3 at the maternal-fetal interface in pregnancy. (**A**) UMAP depicting single-cell transcriptomic analysis of the maternal-fetal interface in early gestation. Colors indicate cell type. dS1-3 represent different decidual subsets. For description of other cell populations, see reference 13. (**B**) *ACTA2* is a marker gene for dS1 cells whereas *DCN* is expressed in both dS2 and dS3 populations. dS3 cells are further characterised by expression of specific decidual genes (e.g. *IL1RL1*, *PRL*, *CLEC2D,* and *CD82*) and enhanced expression of genes involved in glycolysis (e.g. *GAPDH*) and mitochondrial functions (e.g. *NDUFB2*). Genes were visualized using the on-line web tool: https://maternal-fetal-interface.cellgeni.sanger.ac.uk/.
